# YAMP: a containerised workflow enabling reproducibility in metagenomics research

**DOI:** 10.1101/223016

**Authors:** Alessia Visconti, Tiphaine C. Martin, Mario Falchi

## Abstract

YAMP is a user-friendly workflow that enables the analysis of whole shotgun metagenomic data while using containerisation to ensure computational reproducibility and facilitate collaborative research. YAMP can be executed on any UNIX-like system, and offers seamless support for multiple job schedulers as well as for Amazon AWS cloud. Although YAMP has been developed to be ready-to-use by non-experts, bioinformaticians will appreciate its flexibility, modularisation, and simple customisation.

The YAMP script, parameters, and documentation are available at https://github.com/alesssia/YAMP.

## 1 Background

Thanks to the increased cost-effectiveness of high-throughput technologies, the number of studies collecting and analysing large amounts of data has surged, opening new challenges for data analysis and research reproducibility. A ubiquitous lack of repeatability and reproducibility has in fact been observed, and a recent Nature’s survey of 1,576 researchers showed that more than 50% and 70% of them failed to reproduce their own and other scientists’ experiments, respectively [1]. Unavailability of primary data and computational experimentation have been named as the major culprits for this reproducibility crisis, with many studies relying on *ad hoc* scripts and not publishing the necessary code and/nor sufficient details to reproduce the reported results [2,3,4], and with variations across workstations and operating systems representing another obstacle [5,6]. To overcome this issue, tools allowing the development of workflows [7] and software containers [8] have been proposed [9]. In fact, containerised well-structured workflows allow storing every detail of the workflow execution, including software’s versions and parameters (*provenance*, [10]), and nullify systems’ variations [6], guaranteeing studies’ repeatability and reproducibility. Containerised workflows also facilitate collaborative projects, by ensuring identical analysis processes, thus comparable results, and allow the automatisation of data-intensive repetitive tasks [11]. Moreover, they save users with little bioinformatics or computational expertise from the hassles of installing the required pieces of software, and of designing and implementing often complex analysis orchestrations, while expert bioinformaticians can use them as a starting point for customised analyses, thus avoiding redundant solutions.

In metagenomic research, several analysis pipelines have been developed so far. However, they either do not support containerisation (*e.g*., MetAMOS [12], MOCAT2 [13], RAMMCAP [14]), thus potentially compromising reproducibility, or require users to upload their unpublished and/or confidential data on third-party servers (*e.g*., IMG/M [15], the EBI metagenomic pipeline [16], and MG-RAST [17]), where, according to the available resources, they can spend several days waiting to be processed [18], and with data privacy concerning some of the researchers [19]. Scalable metagenomics pipelines allowing both local and cloud execution have been proposed, such as CloVR-Metagenomics [20], and those implemented using the Galaxy platform [21,22]. However, the former lacks steps for quality control (QC) and allows processing reads only generated with the Roche 454 pyrosequencing platform [23], and the latter requires nontrivial expertise for local installation [24], with porting issues observed among different Galaxy versions [25]. QC is also often overlooked. For instance, both MetAMOS and the EBI metagenomic pipeline do not include a step for removing contaminant genomes, with the latter also not discarding identical duplicates. Ignoring decontamination may lead to reads not belonging to the studied ecosystem to be used in downstream analyses, causing potential mismapping on reference databases and, thus, erroneous functional profiling, especially in low biomass environments [26]. Moreover, the presence of contaminating human reads raises privacy concerns, with already two studies able to mine and exploit hosts’ genetic material from publicly available metagenomic samples [26,27]. Retaining duplicated reads, usually considered as technical artefacts derived from PCR amplification [28], may hamper the correct estimation of both community composition and functional capabilities [29]. Finally, MG-RAST performs de-duplication after quality trimming, potentially introducing biases due to the fact that trimming, by modifying the read sequence, may mask true duplicates or generate false ones.

Here we present *“Yet Another Metagenomic Pipeline”* (YAMP), a ready-to-use containerised workflow that, using state-of-the-art tools, processes raw shotgun metagenomic sequencing data up to the taxonomic and functional annotation (Scicrunch.org RRID: SCR_016236). YAMP is implemented in Nextflow [6] and it is accompanied by a Docker and a Singularity [30] container.

## 2 The YAMP workflow

The YAMP workflow is composed of three analysis blocks: the quality control, (**Figure 1**, green rectangle), complemented by several steps of assessment and visualisation of data quality (**Figure 1**, orange rectangle), and the community characterisation (**Figure 1**, pink rectangle).

**Fig. 1.**
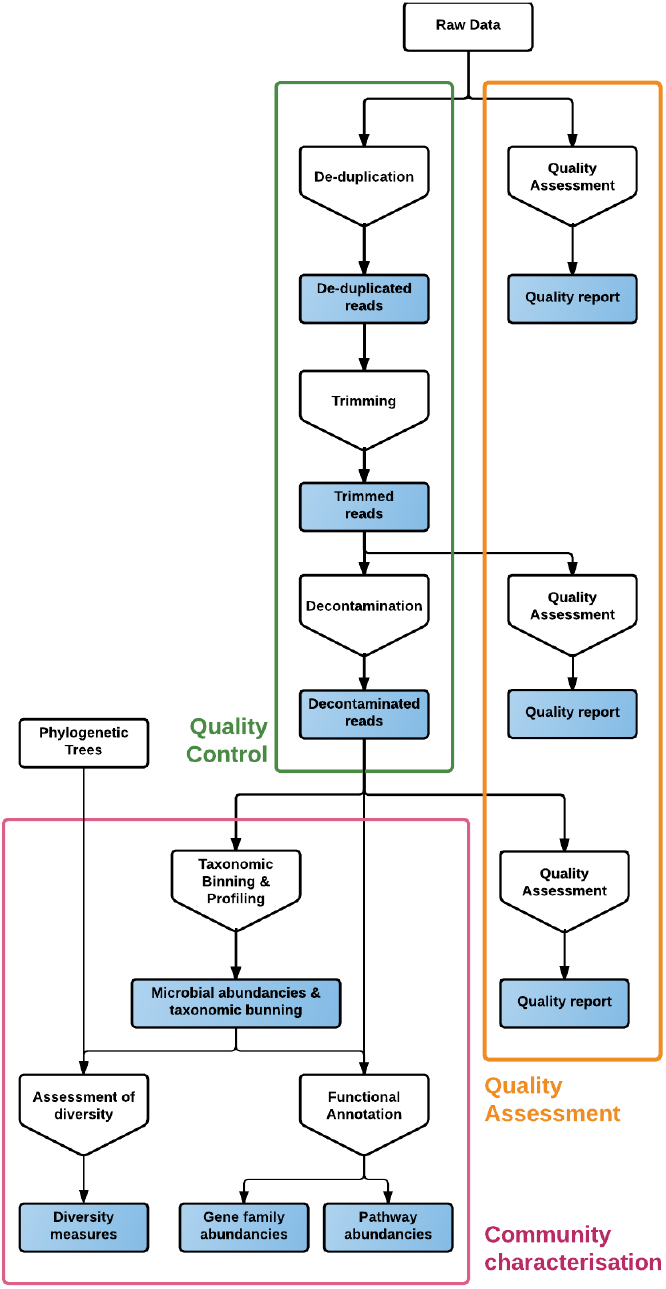
The YAMP workflow. White rectangles represent data to be provided as input, and blue rectangles those produced in output. Pentagons represent the analysis steps.

The QC starts with an optional step of de-duplication, where identical reads, potentially generated by PCR amplification, are removed. The optionality of this step allows retaining *natural duplicates* when PCR-free library preparation approaches (*e.g*., TruSeq) are used. Next, reads are first filtered to remove adapters, known artefacts, phiX, and then quality-trimmed. Notably, YAMP removes duplicates before trimming, avoiding the introduction of biases due to the reads’ sequence modifications. Reads that become too short after trimming are discarded. Indeed, they may map to multiple genomes or genomic regions and compromise downstream analyses. When paired-end reads are at hand, singleton reads (*i.e*., paired-end reads whose mates have been removed) are preserved in order to retain as much information as possible. Finally, reads are screened for contaminants, *e.g*., reads that do not belong to the studied ecosystem. It should be kept in mind, when preparing the custom database of contaminant reads, that many low-complexity sequences and certain features (*e.g*., ribosomes) are highly conserved among species and should be eventually removed to avoid false positive matches.

The implemented QC is accompanied by multiple steps which assess and visualise the reads’ quality, in order to evaluate the quality of the raw data and effectiveness of the trimming and decontamination step.

The QC is followed by multiple steps aimed at estimating multiple *α*-diversity measures, and at characterising the taxonomic and functional profiles of the microbial community, *i.e*. at identifying and quantifying the micro-organisms present in the metagenomic sample (taxonomic binning and profiling) and their functional capabilities (functional characterisation).

## 3 Implementation

YAMP is developed in Nextflow, a workflow management system that allows the effortlessly developing, deploying, and executing of complex distributed computational workflows [6], and which has been used in several life-science projects (*e.g*., [31,32,33]). Nextflow allows for user-transparent high-level parallelisation, and offers out-of-the-box support for distributed computational environments, ensuring the scalability of large projects. Its executor allows porting of workflows on any UNIX-based system (*e.g*., local machine, HPC facilities) in a seamless fashion. Reproducibility is guaranteed by a user-transparent integration with Docker/Singularity and with the BitBucket (https://bitbucket.org/), GitHub (https://github.com/), and GitLab (https://about.gitlab.com/) code repositories, therefore ensuring a consistent tracking of both software and code version. The so-called *retrospective provenance, i.e*., the description of each completed analysis step along with details about its underlying execution environment [10], is captured by task execution reports, which record, among the others, the exact command executed, the tasks’ working directory, environment and output, as well as the container image.

YAMP is accompanied by a Docker and a Singularity [30] container. While Docker defines a platform-independent virtualised light-weight operating system that includes all the pieces of software required by YAMP and traces their versioning, Singularity allows these features to be transferred to High Performance Computing (HPC) systems, with which Docker is inherently incompatible. Along with this single-container approach, YAMP also supports a multi-container scenario. Indeed, while the former is of easier management for users with limited computational experience and allows a more agile deployment, the latter accommodates for YAMP customisation without losing the advantages of a containerised solution in terms of reproducibility and ease of setup.

YAMP integrates state-of-the-art tools for the analysis of metagenomic data. The QC is performed by means of a number of tools belonging to the BBmap suite [34], namely clumpify, BBduk, and BBwrap, which are well-established, and allow processing both single- and paired-end reads from all the major sequencing platforms (*i.e*., Illumina, Roche 454 pyrosequencing, Sanger, Ion Torrent, PacBio, and Oxford Nanopore). They are also computationally efficient, thus scalable to large metagenomic projects and samples. FastQC [35], which provides very detailed reports on reads’ quality, is used to perform QC assessment and visualisation. Taxonomic binning and profiling is performed with MetaPhlAn2 [36], which uses clade-specific markers to both detect the micro-organisms and to estimate their relative abundance. The clade-based approach implemented in MetaPhlAn2 has been shown to scale to large datasets, and to be effective to quantitatively profile the microbial composition during the Human Microbiome Project (HMP) [37] and the Critical Assessment of Metagenome Interpretation (CAMI) challenge [38]. Moreover, MetaPhlAn2 is becoming the *de facto* standard when samples belong to well-characterised environments [39]. The functional capabilities of the microbial community are assessed by the HUMAnN2 pipeline [40], an extension of the pipeline originally developed by the HMP Metabolic Reconstruction Working Group to infer the functional and metabolic potential of microbial communities during the HMP [41]. Briefly, the HUMAnN2 pipeline first stratifies the community in known and unclassified organisms using the MetaPhlAn2 results and the ChocoPhlAn pan-genome database, and then combines these results with those obtained through an organism-agnostic search on the UniRef proteomic database, and the MetaCyc database of metabolic pathways and enzymes [42]. The identified taxonomic profile is additionally used by YAMP to evaluate multiple *α*-diversity measures through the alpha_diversity.py function available in the widely-used QIIME pipeline [43]. QIIME is an extremely modular and efficient pipeline designed for the analysis of amplicon (*e.g*., 16S or 18S rRNA genes) sequencing data, which implements a number of functions investigating ecological features relevant also in metagenomics research.

## 4 YAMP Input/Output

YAMP accepts in input both single- and paired-end FASTQ files, and users can customise the workflow execution either by using command line options or by modifying a simple plain-text configuration file, where parameters are set as key-value pairs. While the parameters should be tuned according to the dataset at hand, to facilitate non-expert users in their analyses of human metagenomic data we provide a set of default parameters derived from our own analysis experience. More in details, we suggest retaining bases with a Phred score of at least 10 (Q10), representing a base call accuracy of 90%, *i.e*., the probability of calling a base out of ten incorrectly. This allows the retrieval also of low coverage regions, therefore improving the total genome recovery and contiguity, an aspect of utmost importance when the QC reads are used for assembly. We also recommend conservatively discarding all reads shorter than 60 bp after trimming, corresponding to a complexity of 4^60^ or less. This length is considered the lowest for avoiding spurious signals when carrying on functional characterisation via HUMAnN2 [44,45,46]. Next, we propose using a minimum alignment identity of 95% (maximum indel length: 3 bp) to identify contaminants reads, which has been shown to possibly zero the number of false positive when used on opportunely created custom database of contaminant reads [47]. Finally, we suggest using the UniRef90 protein database, as its clusters are more likely to be isofunctional and non-redundant. However, the UniRef50 protein database should be preferred when dealing with poorly characterised microbiomes.

The output generated by YAMP includes a FASTQ file of QC’ed reads, the taxonomy composition along with the microbe, gene and pathway relative abundances, the pathway coverage, and multiple *α*-diversity measures. An option allows users to retain temporary files, such as these generated by the QC steps or during the HUMAnN2 execution. Additionally, YAMP outputs several QC reports, a very detailed log file recording information about each analysis step which ensures the retrospective provenance (**Figure 2**), and statistics of memory usage and time of execution (**Figure 3**). It should be noted that the disk space requested by the files generated by YAMP is, in average, about seven times the size of the raw data files, and particular attention should be paid when multiple samples are processed simultaneously, as in multicore machines or HPC facilities. However, discarding temporary files, as we suggest, will require a final disk space of 20-70% the size of the raw data files, with higher quality files requiring more space due to the small number of reads discarded during the QC process.

**Fig. 2.**
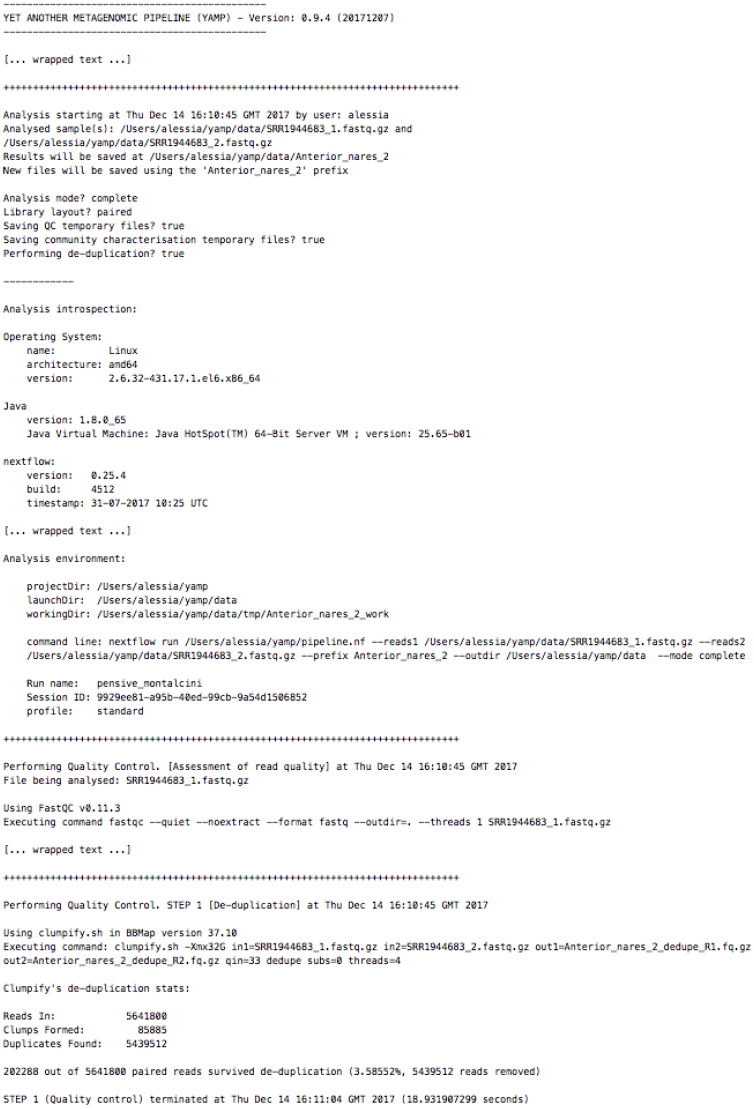
Example of an excerpt of the YAMP execution log.

**Fig. 3.**
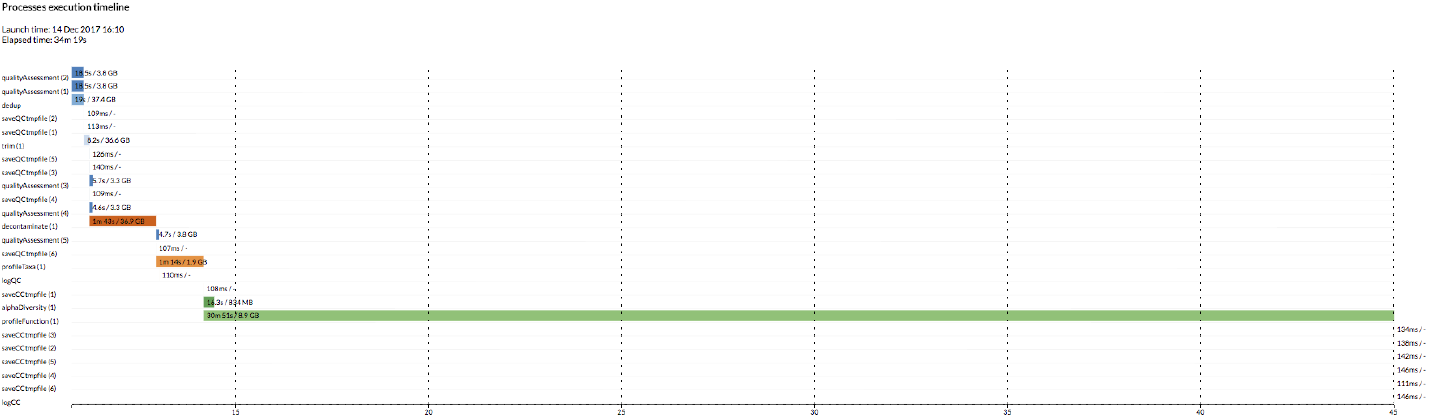
Example of YAMP execution profile. YAMP returns the time spend during its complete execution and in each step, as well as the steps’ memory peaks.

## 5 Results

To compare YAMP to existing metagenomics analysis workflows we simulated datasets including either different percentages of human contamination or of artificial duplicates. Then, to facilitate the discussion on YAMP computational requirements, and to assess its ability to reproduce research results already described in the literature, we carried out a real-word case study, which included 18 samples collected in different body sites. Notably, despite both the simulation and the real word case study focus on human metagenomics data, YAMP can be used for the analysis of data originated from virtually any environment.

### 5.1 Simulation study

In the first simulated scenario, we aimed at testing the impact of contaminant reads on the functional characterisation of the microbial community. Therefore, we generated five metagenomics samples simulating a human oral community with 13 bacterial species, and including a variable amount of human reads as contaminants. The relative proportion of the bacterial genomes followed that suggested by Zhou *et al*. [48] (**Supplementary Table S1**). The percentage of contaminant reads was of 1, 5, 25, 50, and 80%, in line with the amounts observed in the literature for human samples [26,49,50]. For instance, the HMP Consortium targeted as human 49% of the total reads, also observing that samples collected from soft tissue and preparations from saliva showed the highest human contamination, with samples from mid-vagina, saliva, anterior nares, and throat including 96, 80, 82, and 75% of human DNA sequence, respectively, whereas stool was only marginally affected, including less than 1% of human contamination [50]. We compared the functional profiles inferred by YAMP with those generated using the EBI metagenomics pipeline (version 4.1), which does not include a decontamination step. These two workflows use different databases for the functional characterisation (*i.e*., MetaCyc, and both the InterPro and the Gene Ontology databases, respectively), and their results could not be directly compared. Therefore, for both tools, we evaluated the root-mean-square error (RMSE) between the functional profiles inferred for each of the contaminated dataset and that inferred for a baseline dataset with no human contamination (the lowest the RMSE, the better the fit). YAMP showed the best performances, with a RMSE < 1.25 × 10^−6^ regardless of the amount of human contamination (**Table 1, Supplementary Figure S1**). Notably, despite YAMP was not able to remove all the contaminants reads (likely due to the masking of low-complexity and highly conserved region in the reference genome used during the decontamination step), its performances are stable, plausibly thanks to the high specificity of the annotation database used. The EBI metagenomics pipeline guaranteed appreciable results: the maximum RMSE was 0.086 with 80% human contamination and using functional annotations from the Gene Ontology (GO) Slim database. However, while its GO-based results showed errors that were uniformly distributed along the identified annotations, and which increased with the level of human contamination, results on the InterPro database seemed to be concentrate on few specific domains (**Supplementary Figure S1**). Three of these domains (*“L1 transposable element, dsRBD-like domain”, “L1 transposable element, trimerization domain”, “Domain of unknown function DUF1725”*), which were not detected in the baseline dataset, are connected to the long interspersed nuclear element 1 (LINE-1 or L1), an active retrotransposon, which comprises approximately 17% of the human genome [51]: an obvious sign of uncontrolled contamination.

**Table 1.**
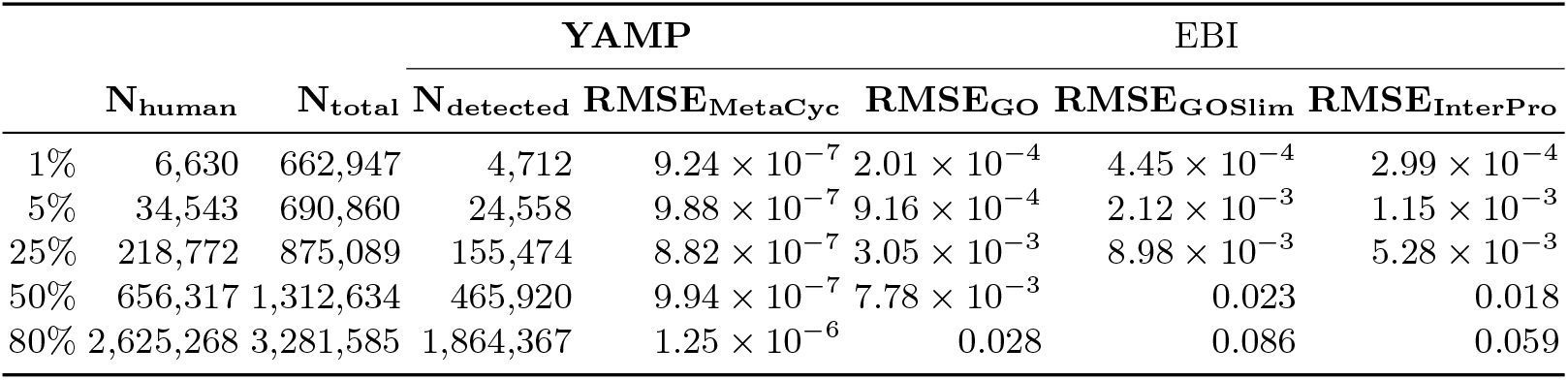
Results of the first simulation study (human contamination). For each level of human contamination, we report the number of human and total reads, and, only for YAMP, the number of human reads correctly detected and removed. *RMSE* values were evaluated on the inferred proportions using a dataset with no human contamination as a baseline. For the EBI metagenomics pipeline (v4.1), we report RMSE values evaluated on the three databases used for functional characterisation.

In the second simulated scenario, we aimed at testing the impact of artificial duplicates on the microbial community’s functional characterisation. Therefore, we generated three datasets simulating the human oral community described before (**Supplementary Table S1**) but without any natural duplicates, and introducing percentages of duplication of about 0.25, 1.25, and 5%, in line with the values described in the literature [52,53]. Consistently with the observations that GC-rich DNA sequences are difficult to amplify, and that the lower the CG content the higher the probability of an amplification bias [52], we allowed the introduction of duplicates mostly from bacteria with the lowest GC-content, namely *Streptococcus peroris, Veillonella atypica, Veillonella parvula* and *Veillonella dispair* (**Methods**). We compared the functional profiles generated by YAMP with both those generated using the EBI metagenomics pipeline (version 4.1), which does not include a de-duplication step, and those generated using the MG-RAST pipeline (version 4.0.3), which performs trimming before de-duplication, potentially introducing biases. As in the previous simulation study, we assessed their performances on a baseline dataset that in this case did not included any duplicate. YAMP was again the best performer, with a RMSE < 1.17 × 10^−6^ regardless of the amount of duplication, and with stable results at each duplication level (**Table 2, Supplementary Figure S2**). The EBI metagenomics pipeline offered excellent results when evaluated over the InterPro database annotation (RMSE < 6.85 × 10^−3^), but its performance slightly degraded when the evaluation was performed on the GO and GO Slim annotations, where we observed, at a 5% level of duplication, a maximum RMSE of 0.011 and 0.022, respectively (**Table 2**). MG-RAST allows functional characterisation to be performed using several databases (*e.g*., RefSeq, GenBank, SEED, KEGG, …). For the sake of simplicity, here we used the annotations derived from the SEED Subsystems ontology [54] (which are provided as a pre-computed summary of the reads assigned at the highest level of this functional hierarchy), and from the KEGG database (which we preprocessed to extract the number of reads assigned at each KEGG annotation – level 3; see **Methods**). MG-RAST performances were acceptable when evaluated on the KEGG database (RMSE < 0.017), but decreased sensibly on the SEED Subsystems ontology (RMSE > 0.243, **Table 2**), mostly due to a progressive over-estimation of the *“Nucleosides and Nucleotides”* ontology term (estimated to be 3.69, 4.93, 4.98, 5.27% at baseline and at 0.25, 1.25, and 5% percentage of duplication, respectively).

**Table 2.**
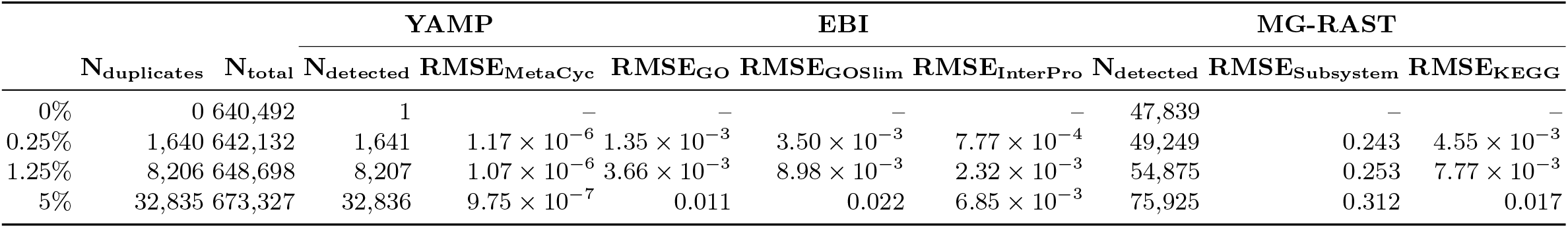
Results of the second simulation study (artificial duplicates). For each level of artificial duplicates, we report the number of duplicates and total reads, and, for YAMP and MG-RAST, the number of duplicated reads removed. *RMSE* values were evaluated on the inferred proportions using a dataset with no duplicated reads as a baseline. For the EBI metagenomics pipeline (v4.1), we report RMSE values evaluated on the three databases used for functional characterisation; for MG-RAST (v4.0.3), we report RMSE values evaluated on two of the databases available for functional characterisation (*i.e*., SEED Subsystem and KEGG).

### 5.2 Real word case study

We analysed 18 randomly selected samples from six different body sites sequenced during the Phase III of the Human Microbiome Project [50] (**Table 3**). On average, the selected samples included 12.6M paired-end reads (25.2M reads in total), which yielded to 13.3M QC’ed reads (including both paired-end and singleton reads), and were processed in an average time of two hours using four threads on a machine sporting a 2.60GHz Intel^®^ Xeon^®^ processor with 32 GB of RAM and using the default YAMP parameters (**Table 3**). At the phylum level, each body site showed a characteristic signature (**Figure 4**), with a predominance of Actinobacteria in the airways, Firmicutes in the vagina, Bacteroidetes in the stool, and a mixture of Actinobacteria, Firmicutes and Proteobacteria in the oral cavity, as already observed in previous studies [55]. A site-specific microbial signature was also present at the species level, where both the Principal coordinate analysis (PCoA) evaluated using the Bray-Curtis dissimilarity (**Supplementary Figure S3 and S4**), and the hierarchical clustering computed on the Manhattan distances between species relative abundances (**Supplementary Figure S5**) showed that the taxonomy composition was sufficient to discriminate among body sites, even though it had limited ability in distinguishing among different loci in the oral cavity.

**Fig. 4.**
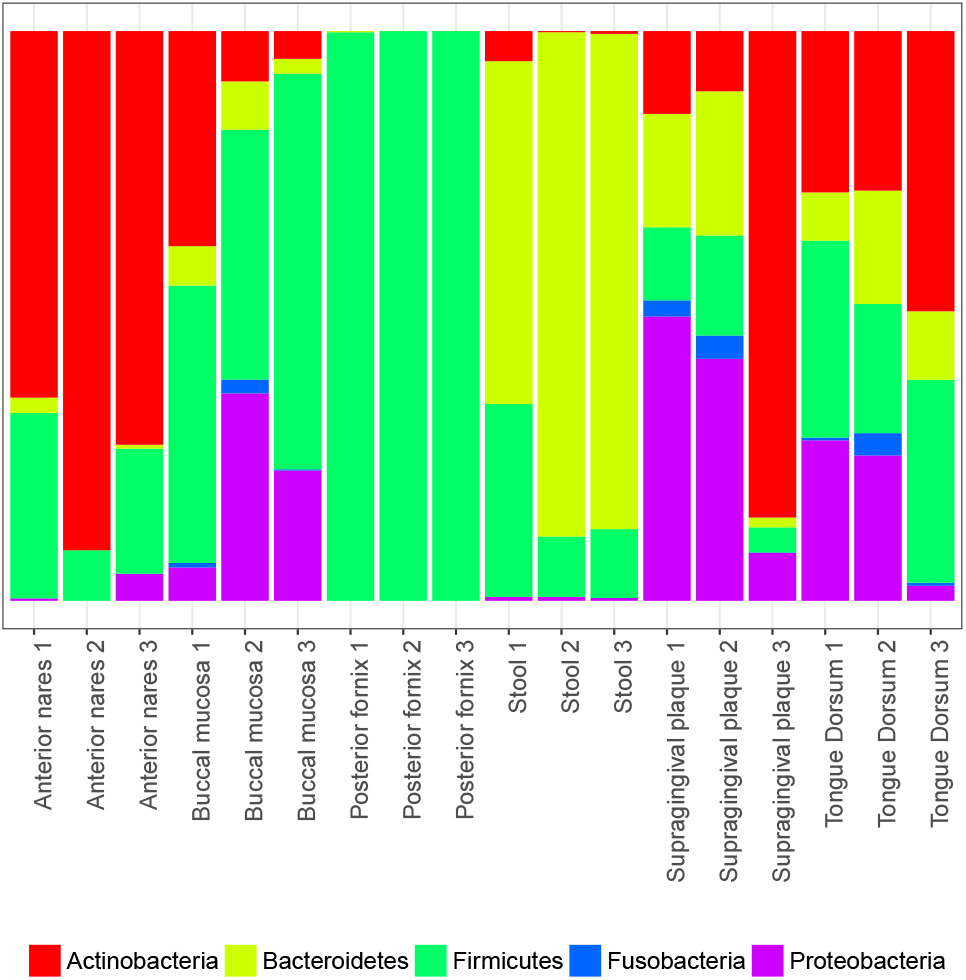
Phylum level relative abundances. Each vertical bar represents a sample. Phylum relative abundances were estimated by YAMP using MetaPhlAn2. Unspecified viral phyla are not shown.

**Table 3.**
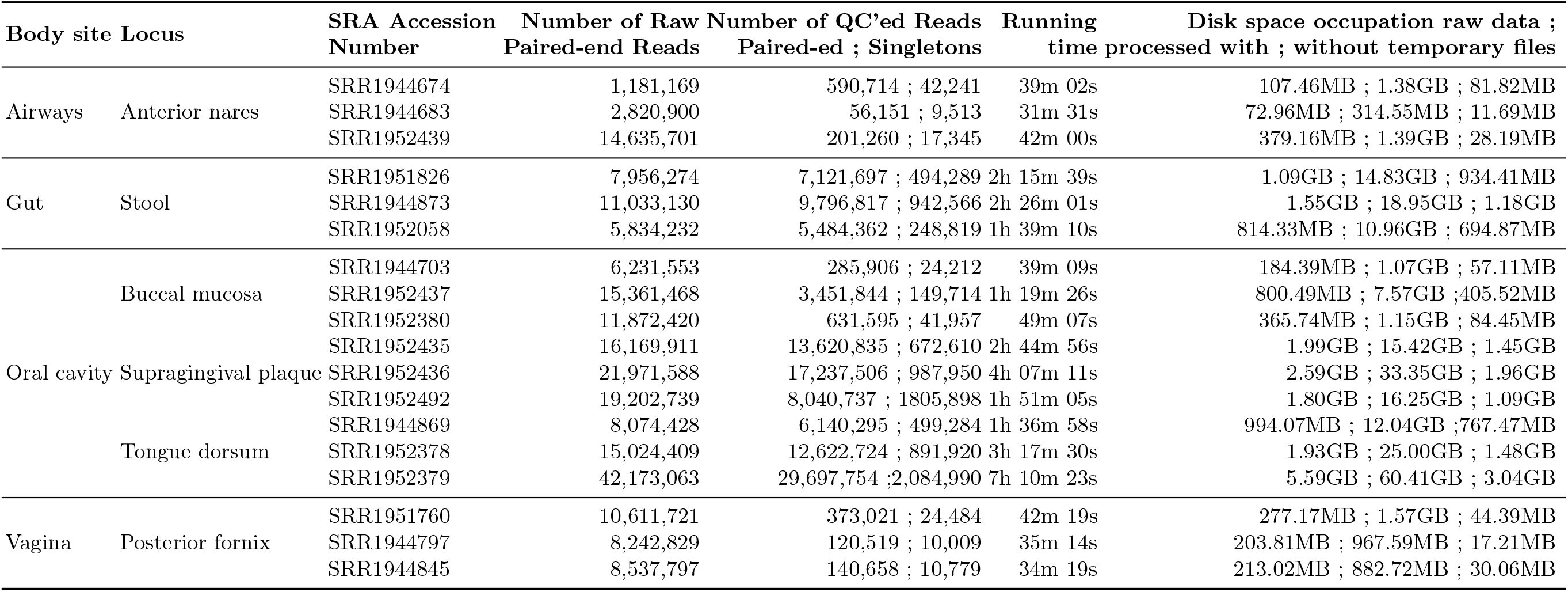
Run Accession Number and statistics for **18** randomly selected samples from the Human Microbiome Project (HMP) Phase III [50]. Samples were processed using 4 threads on a machine sporting a 2.60GHz Intel^®^ Xeon^®^ processor with 32 GB of RAM. The required disk space does not include the size of the raw data file.

## 6 Discussion

In conclusion, with YAMP, we provide a user-friendly workflow that enables the analysis of whole shotgun metagenomic data. By supporting containerisation, YAMP allows for computational reproducibility, also enabling collaborative studies. In fact, while software versions are described in the Docker/Singularity container, the Nextflow script and configuration file capture all the details needed to fully track each step of data processing, thus satisfying the prospective provenance requirements, while the very detailed YAMP log file ensures retrospective provenance. Indeed, to ensure reproducibility, researchers should only provide the YAMP configuration file and a link to the container image. Being based on Nextflow, YAMP runs on any UNIX-like system, provides out-of-the-box support for several job schedulers (*e.g*., PBS, SGE, SLURM) and for Amazon AWS cloud, and its integration with Docker/Singularity is completely user-transparent. Finally, while YAMP has been developed to be ready to use by non-experts, and potentially does not require any software installation or parameter tuning, bioinformaticians will value its flexibility and simple customisation. In fact, the well-defined YAMP modularisation and the usage of standard data formats allow both an easy integration of new analysis steps and a customisation of the existing ones. This is of particular importance in a fast-developing field such as metagenomics, where the analysis guidelines and tools and are not stable yet. With YAMP we provide a workflow to which new analysis modules can be easily added, and where tools that become outdated can be effortlessly replaced, therefore securing its sustainability.

YAMP is made available as a Nextflow script that allows a user-friendly execution via the command line. The source code is available in the YAMP GitHub repository (https://github.com/alesssia/YAMP), which includes a wiki with a full documentation and several tutorials (https://github.com/alesssia/YAMP/wiki). The Docker/Singularity image can be downloaded and installed from DockerHub (https://hub.docker.com/r/alesssia/yampdocker).

## 7 Potential implications

YAMP has been designed with the specific goals of enabling reproducible metagenomics analyses, facilitating collaborative projects, and helping researchers with limited computational experience who are approaching this field of research. However, we are confident that other areas of research would be aided by a more widespread use of containerised well-structured workflows. Indeed, as outlined in the Background Section, a lack of reproducibility is nowadays ubiquitous, and, besides undermining the credibility of scientific research, it has an economical cost, quantified, for instance, in US$28B/year for preclinical research [56]. On the other hand, ensuring reproducibility does not come for free: anecdotic evidence suggests that the time spent on a project may increase by 30-50% [1], and that to reproduce the analysis of single computational biology paper can require up to 280 hours [57]. YAMP, along with other containarised workflows, such as the Integrated Meta-omic Pipeline (IMP) [58] and Bio-Docklets [59], represents a proof-of-concept showing a simple way to enable reproducible and collaborative research. We also advocate the sharing of such containerised workflows, which will benefit a wide group of researchers, regardless of their computational experience [11].

## 8 Methods

### 8.1 Data Availability

#### Simulation Study

The simulated metagenomic samples were generated using the reference human and microbial genomes downloaded from NCBI (http://www.ncbi.nlm.nih.gov; **Supplementary Table S1**, human genome: build GRCh37.p13). Each microbial genome was processed independently, and used as input for the randomreads tool from the BBmap suite [34], which allows generating random reads in various formats. Oral bacterial single-end sequences were generated using the default parameters, and asking for a fixed read length of 100bp, a Phred score ranging from 6 to 40 (average: 20), and with a probability of mutation (SNP, insertion, deletion, substitution, and N calls) of 0.2 (max 10 blocks of Ns per read). We also assigned scaffolds a random exponential coverage level, to simulate a metagenomic coverage distribution (metagenome=t), and asked for a fixed genomic coverage (**Supplementary Table S1**). Human singleend sequences were created using the same parameters used for the microbial genomes and asking for a fixed number of reads (**Table 1**).

In the second simulation scenario, we first de-duplicated each of the microbial simulated datasets generated beforehand using clumpify [34], and, for each of the four species showing the lowest GC-content (*Streptococcus peroris, Veillonella atypica, parvula and dispair*), we generated sets of duplicated reads by randomly selecting and matching reads and quality scores in order to include 40% of identical duplicates. Then, we merged the de-duplicated dataset with these sets of duplicated reads to build a new reads pool. Next, we randomly selected, from the described pool, fixed numbers of reads which were finally merged with the de-duplicated dataset in order to generate each simulated sample. This allowed duplicates more likely to belong but not limited to the GC-poor microbial genomes, and to have duplicated reads which are more likely but not limited to have different quality scores.

The simulated datasets are available from the European Nucleotide Archive website (Study accession numbers: PRJEB25791 and PRJEB26333; **Supplementary Table S2**).

#### Real word case study

The 18 randomly selected samples used to assess YAMP belong to the Phase III of the Human Microbiome Project [50], and were downloaded from the European Nucleotide Archive website (Study accession number: PRJNA275349). Samples were collected from healthy adults residing in the USA at the time of sample collection. After genomic DNA extraction, metagenomics library preparation was performed using the NexteraXT library construction protocol. Paired-end metagenomic sequencing was performed on the Illumina HiSeq2000 platform with a read length of 100 bp. Samples’ accession numbers are reported in **Table 3**.

#### 8.2 Data Analysis

Simulation Study Samples were processed with YAMP using the default parameters, as defined in the published YAMP configuration file (https://raw.githubusercontent.com/alesssia/YAMP/master/nextflow.config), and the databases deposited on Zenodo (https://doi.org/10.5281/zenodo.1068229). Unmapped reads were discarded. When analysing the simulated dataset with the EBI metagenomics pipeline (version 4.1) and MG-RAST (version 4.0.3), the default settings were used. Results from the EBI metagenomics pipeline were downloaded via the web user interface, and the functional profiles were evaluated by transforming to percentage the proportion of reads assigned to each function. When evaluating MG-RAST performances, we used the functional profiles evaluated on the SEED Subsystems ontology and KEGG database, identified at a default alignment length > 15bp, e-value < 1 × 10^−5^, and percent identity > 60% (as by MG-RAST default). Data for the SEED Subsystems ontology, annotated at the highest level of the hierarchy, were downloaded via the web user interface, and the functional profiles were evaluated by transforming to percentage the proportion of reads assigned to each function. Data for the KEGG database were downloaded via the web user interface, and pre-processed to extract the number of reads assigned to each KEGG function (level 3; reads assigned to multiple annotations were discarded), and then transformed to percentage as explained beforehand.

#### Real word case study

Samples were processed with YAMP using the default parameters, as defined in the published YAMP configuration file (https://raw.githubusercontent.com/alesssia/YAMP/master/nextflow.config), and the databases deposited on Zenodo (https://doi.org/10.5281/zenodo.1068229). The Bray-Curtis dissimilarity values were evaluated using the species relative abundances as estimated by YAMP using MetaPhlAn2 [36] and the *vegdist* function in the vegan R package (version 2.4.3) [60]. Principal coordinate analysis (PCoA) was evaluated on the Bray-Curtis dissimilarity values using the *pcoa* function in the ape R package (version 4.1) [61]. Hierarchical clustering was computed using the Manhattan distance between species relative abundances and the *pvclust* function in the pvclust R package (version 2.0) [62]. 10,000 bootstrap interactions were used to evaluate the P values supporting each cluster.

## 9 Availability of source code and requirements

– Project name: YAMP
– Project home page: https://github.com/alesssia/YAMP
– Operating system(s): UNIX-like systems, support for Amazon AWS Cloud
– Programming language: Nextflow
– Other requirements: Java, Docker/Singularity
– License: GNU GPL v3
– Any restrictions to use by non-academics: None

## Declarations

### List of abbreviations

HPC: : High Performance Computing;
PCoA: : Principal coordinate analysis;
QC: : Quality Control;
RMSE: : root-mean-square error.

## Author’s Contributions

AV, TCM, and MF designed the metagenomics workflow. AV implemented and optimised the workflow, created the Docker container, and wrote the manuscript. All the authors read, commented, and approved the final manuscript.

## Acknowledgements

AV would like to thank Paolo Di Tommaso and Brian Bushnell for their help with Nextflow and BBmap.

## References

1. M. Baker, “1,500 scientists lift the lid on reproducibility,” Nature News, vol. 533, no. 7604, p. 452, 2016.

2. J. P. Ioannidis, D. B. Allison, C. A. Ball, I. Coulibaly, X. Cui, A. C. Culhane, M. Falchi, C. Furlanello, L. Game, G. Jurman, et al., “Repeatability of published microarray gene expression analyses,” Nature genetics, vol. 41, no. 2, pp. 149–155, 2009.

3. T. Hothorn and F. Leisch, “Case studies in reproducibility,” Briefings in bioinformatics, vol. 12, no. 3, pp. 288–300, 2011.

4. R. D. Peng, “Reproducible research in computational science,” Science, vol. 334, no. 6060, pp. 1226–1227, 2011.

5. E. H. Gronenschild, P. Habets, H. I. Jacobs, R. Mengelers, N. Rozendaal, J. Van Os, and M. Marcelis, “The effects of FreeSurfer version, workstation type, and Macintosh operating system version on anatomical volume and cortical thickness measurements,” PloS one, vol. 7, no. 6, p. e38234, 2012.

6. P. Di Tommaso, M. Chatzou, E. W. Floden, P. P. Barja, E. Palumbo, and C. Notredame, “Nextflow enables reproducible computational workflows,” Nature Biotechnology, vol. 35, no. 4, pp. 316–319, 2017.

7. J. Leipzig, “A review of bioinformatic pipeline frameworks,” Briefings in bioinformatics, vol. 18, no. 3, pp. 530–536, 2017.

8. C. Boettiger, “An introduction to Docker for reproducible research,” ACM SIGOPS Operating Systems Review, vol. 49, no. 1, pp. 71–79, 2015.

9. S. R. Piccolo and M. B. Frampton, “Tools and techniques for computational reproducibility,” GigaScience, vol. 5, no. 1, p. 30, 2016.

10. S. B. Davidson and J. Freire, “Provenance and scientific workflows: challenges and opportunities,” in Proceedings of the 2008 ACM SIGMOD international conference on Management of data, pp. 1345–1350, ACM, 2008.

11. O. Spjuth, E. Bongcam-Rudloff, G. C. Hernández, L. Forer, M. Giovacchini, R. V. Guimera, A. Kallio, E. Korpelainen, M. M. Kańduła, M. Krachunov, et al., “Experiences with workflows for automating data-intensive bioinformatics,” Biology direct, vol. 10, no. 1, p. 43, 2015.

12. T. J. Treangen, S. Koren, D. D. Sommer, B. Liu, I. Astrovskaya, B. Ondov, A. E. Darling, A. M. Phillippy, and M. Pop, “MetAMOS: a modular and open source metagenomic assembly and analysis pipeline,” Genome biology, vol. 14, no. 1, p. R2, 2013.

13. J. R. Kultima, L. P. Coelho, K. Forslund, J. Huerta-Cepas, S. S. Li, M. Driessen, A. Y. Voigt, G. Zeller, S. Sunagawa, and P. Bork, “MOCAT2: a metagenomic assembly, annotation and profiling framework,” Bioinformatics, vol. 32, no. 16, pp. 2520–2523, 2016.

14. W. Li, “Analysis and comparison of very large metagenomes with fast clustering and functional annotation,” BMC bioinformatics, vol. 10, no. 1, p. 359, 2009.

15. V. M. Markowitz, I.-M. A. Chen, K. Chu, E. Szeto, K. Palaniappan, M. Pillay, A. Ratner, J. Huang, I. Pagani, S. Tringe, et al., “IMG/M 4 version of the integrated metagenome comparative analysis system,” Nucleic Acids Research, vol. 42, no. D1, pp. D568–D573, 2013.

16. A. L. Mitchell, M. Scheremetjew, H. Denise, S. Potter, A. Tarkowska, M. Qureshi, G. A. Salazar, S. Pesseat, M. A. Boland, F. M. Hunter, et al., “EBI Metagenomics in 2017: enriching the analysis of microbial communities, from sequence reads to assemblies,” Nucleic acids research, 2017.

17. F. Meyer, D. Paarmann, M. D’Souza, R. Olson, E. M. Glass, M. Kubal, T. Paczian, A. Rodriguez, R. Stevens, A. Wilke, et al., “The metagenomics RAST server– a public resource for the automatic phylogenetic and functional analysis of metagenomes,” BMC bioinformatics, vol. 9, no. 1, p. 386, 2008.

18. A. Wilke, W. Gerlach, T. Harrison, T. Paczian, W. L. Trimble, and F. Meyer, “MGRAST Manual for version 4, revision 3.” ftp://ftp.metagenomics.anl.gov/data/manual/mg-rast-tech-report-v4-r3.pdf 2017.

19. E. Pérez-Wohlfeil, J. A. Arjona-Medina, O. Torreno, E. Ulzurrun, and O. Trelles, “Computational workflow for the fine-grained analysis of metagenomic samples,” BMC genomics, vol. 17, no. 8, p. 802, 2016.

20. S. V. Angiuoli, M. Matalka, A. Gussman, K. Galens, M. Vangala, D. R. Riley, C. Arze, J. R. White, O. White, and W. F. Fricke, “CloVR: a virtual machine for automated and portable sequence analysis from the desktop using cloud computing,” BMC bioinformatics, vol. 12, no. 1, p. 356, 2011.

21. E. Afgan, D. Baker, M. Van den Beek, D. Blankenberg, D. Bouvier, M. Čech, J. Chilton, D. Clements, N. Coraor, C. Eberhard, et al., “The Galaxy platform for accessible, reproducible and collaborative biomedical analyses: 2016 update,” Nucleic acids research, vol. 44, no. W1, pp. W3–W10, 2016.

22. S. K. Pond, S. Wadhawan, F. Chiaromonte, G. Ananda, W.-Y. Chung, J. Taylor, A. Nekrutenko, G. Team, et al., “Windshield splatter analysis with the Galaxy metagenomic pipeline,” Genome research, vol. 19, no. 11, pp. 2144–2153, 2009.

23. J. White, C. Arze, M. Matalka, T. C. Team, O. White, S. Angiuoli, and W. Frickels, “CloVR-Metagenomics: Functional and taxonomic microbial community characterization from metagenomic whole-genome shotgun (WGS) sequences–standard operating procedure, version 1.0,” Nature Preecidings, 2011.

24. E. Ladoukakis, F. N. Kolisis, and A. A. Chatziioannou, “Integrative workflows for metagenomic analysis,” Frontiers in cell and developmental biology, vol. 2, 2014.

25. S. Cohen-Boulakia, K. Belhajjame, O. Collin, J. Chopard, C. Froidevaux, A. Gaignard, K. Hinsen, P. Larmande, Y. Le Bras, F. Lemoine, et al., “Scientific workflows for computational reproducibility in the life sciences: Status, challenges and opportunities,” Future Generation Computer Systems, 2017.

26. S. K. Ames, S. N. Gardner, J. M. Marti, T. R. Slezak, M. B. Gokhale, and J. E. Allen, “Using populations of human and microbial genomes for organism detection in metagenomes,” Genome research, vol. 25, no. 7, pp. 1056–1067, 2015.

27. R. Blekhman, J. K. Goodrich, K. Huang, Q. Sun, R. Bukowski, J. T. Bell, T. D. Spector, A. Keinan, R. E. Ley, D. Gevers, and A. G. Clark, “Host genetic variation impacts microbiome composition across human body sites,” Genome biology, vol. 16, no. 1, p. 191, 2015.

28. H. Xu, X. Luo, J. Qian, X. Pang, J. Song, G. Qian, J. Chen, and S. Chen, “FastUniq: a fast de novo duplicates removal tool for paired short reads,” PloS one, vol. 7, no. 12, p. e52249, 2012.

29. M. B. Jones, S. K. Highlander, E. L. Anderson, W. Li, M. Dayrit, N. Klitgord, M. M. Fabani, V. Seguritan, J. Green, D. T. Pride, S. Yooseph, W. Biggs, K. Nelson, and C. Venter, “Library preparation methodology can influence genomic and functional predictions in human microbiome research,” Proceedings of the National Academy of Sciences, vol. 112, no. 45, pp. 14024–14029, 2015.

30. G. M. Kurtzer, V. Sochat, and M. W. Bauer, “Singularity: Scientific containers for mobility of compute,” PloS one, vol. 12, no. 5, p. e0177459, 2017.

31. C. Guzman and I. Dorso, “CIPHER: a flexible and extensive workflow platform for integrative next-generation sequencing data analysis and genomic regulatory element prediction,” BMC bioinformatics, vol. 18, no. 1, p. 363, 2017.

32. C. L. Cario and J. S. Witte, “Orchid: a novel management, annotation, and machine learning framework for analyzing cancer mutations,” Bioinformatics, 2017.

33. N. D. Sanderson, T. L. Street, D. Foster, J. Swann, B. L. Atkins, A. J. Brent, M. A. McNally, S. Oakley, A. Taylor, T. E. Peto, et al., “Real-time analysis of nanopore-based metagenomic sequencing from orthopaedic device infection,” bioRxiv, p. 220616, 2017.

34. B. Bushnell, “BBMap short-read aligner, and other bioinformatics tools.” https://sourceforge.net/projects/bbmap/, 2015.

35. S. Andrews, “FastQC A Quality Control tool for High Throughput Sequence Data.” http://www.bioinformatics.babraham.ac.uk/projects/fastqc/, 2010.

36. D. T. Truong, E. A. Franzosa, T. L. Tickle, M. Scholz, G. Weingart, E. Pasolli, A. Tett, C. Huttenhower, and N. Segata, “MetaPhlAn2 for enhanced metagenomic taxonomic profiling,” Nature methods, vol. 12, no. 10, p. 902, 2015.

37. H. M. P. Consortium et al., “Structure, function and diversity of the healthy human microbiome,” Nature, vol. 486, no. 7402, pp. 207–214, 2012.

38. A. Sczyrba, P. Hofmann, P. Belmann, D. Koslicki, S. Janssen, J. Droege, I. Gregor, S. Majda, J. Fiedler, E. Dahms, et al., “Critical Assessment of Metagenome Interpretation- a benchmark of computational metagenomics software,” Biorxiv, p. 099127, 2017.

39. C. Quince, A. W. Walker, J. T. Simpson, N. J. Loman, and N. Segata, “Shotgun metagenomics, from sampling to analysis,” Nature Biotechnology, vol. 35, no. 9, pp. 833–844, 2017.

40. S. Abubucker, N. Segata, J. Goll, A. M. Schubert, J. Izard, B. L. Cantarel, B. Rodriguez-Mueller, J. Zucker, M. Thiagarajan, B. Henrissat, et al., “HUMAnN2: The HMP Unified Metabolic Analysis Network 2.” http://huttenhower.sph.harvard.edu/humann2, 2017.

41. S. Abubucker, N. Segata, J. Goll, A. M. Schubert, J. Izard, B. L. Cantarel, B. Rodriguez-Mueller, J. Zucker, M. Thiagarajan, B. Henrissat, et al., “Metabolic reconstruction for metagenomic data and its application to the human microbiome,” PLoS computational biology, vol. 8, no. 6, p. e1002358, 2012.

42. R. Caspi, T. Altman, K. Dreher, C. A. Fulcher, P. Subhraveti, I. M. Keseler, A. Kothari, M. Krummenacker, M. Latendresse, L. A. Mueller, et al., “The metacyc database of metabolic pathways and enzymes and the biocyc collection of pathway/genome databases,” Nucleic acids research, vol. 40, no. D1, pp. D742–D753, 2011.

43. J. G. Caporaso, J. Kuczynski, J. Stombaugh, K. Bittinger, F. D. Bushman, E. K. Costello, N. Fierer, A. G. Peña, J. K. Goodrich, J. I. Gordon, et al., “QIIME allows analysis of high-throughput community sequencing data,” Nature methods, vol. 7, no. 5, pp. 335–336, 2010.

44. M. Schirmer, S. P. Smeekens, H. Vlamakis, M. Jaeger, M. Oosting, E. A. Franzosa, R. ter Horst, T. Jansen, L. Jacobs, M. J. Bonder, et al., “Linking the human gut microbiome to inflammatory cytokine production capacity,” Cell, vol. 167, no. 4, pp. 1125–1136, 2016.

45. B. D. Piening, W. Zhou, K. Contrepois, H. Röst, G. J. G. Urban, T. Mishra, B. M. Hanson, E. J. Bautista, S. Leopold, C. Y. Yeh, et al., “Integrative Personal Omics Profiles during Periods of Weight Gain and Loss,” Cell Systems, 2018.

46. A. F. Schulfer, T. Battaglia, Y. Alvarez, L. Bijnens, V. E. Ruiz, M. Ho, S. Robinson, T. Ward, L. M. Cox, A. B. Rogers, et al., “Intergenerational transfer of antibiotic-perturbed microbiota enhances colitis in susceptible mice,” Nature microbiology, vol. 3, no. 2, p. 234, 2018.

47. B. Bushnell, “Introducing RemoveHuman: Human Contaminant Removal.” http://seqanswers.com/forums/showthread.php?t=42552, 2014.

48. Q. Zhou, X. Su, and K. Ning, “Assessment of quality control approaches for metagenomic data analysis,” Scientific reports, vol. 4, p. 6957, 2014.

49. R. Schmieder and R. Edwards, “Fast identification and removal of sequence contamination from genomic and metagenomic datasets,” PloS one, vol. 6, no. 3, p. e17288, 2011.

50. The Human Microbiome Project Consortium, “A framework for human microbiome research,” Nature, vol. 486, no. 7402, p. 215, 2012.

51. E. Khazina and O. Weichenrieder, “Non-ltr retrotransposons encode noncanonical rrm domains in their first open reading frame,” Proceedings of the National Academy of Sciences, vol. 106, no. 3, pp. 731–736, 2009.

52. V. Gomez-Alvarez, T. K. Teal, and T. M. Schmidt, “Systematic artifacts in metagenomes from complex microbial communities,” The ISME journal, vol. 3, no. 11, p. 1314, 2009.

53. B. Niu, L. Fu, S. Sun, and W. Li, “Artificial and natural duplicates in pyrosequencing reads of metagenomic data,” BMC bioinformatics, vol. 11, no. 1, p. 187, 2010.

54. R. Overbeek, T. Begley, R. M. Butler, J. V. Choudhuri, H.-Y. Chuang, M. Cohoon, V. de Crécy-Lagard, N. Diaz, T. Disz, R. Edwards, et al., “The subsystems approach to genome annotation and its use in the project to annotate 1000 genomes,” Nucleic acids research, vol. 33, no. 17, pp. 5691–5702, 2005.

55. K. Aagaard, J. Ma, K. M. Antony, R. Ganu, J. Petrosino, and J. Versalovic, “The placenta harbors a unique microbiome,” Science translational medicine, vol. 6, no. 237, pp. 237ra65–237ra65, 2014.

56. L. P. Freedman, I. M. Cockburn, and T. S. Simcoe, “The economics of reproducibility in preclinical research,” PLoS biology, vol. 13, no. 6, p. e1002165, 2015.

57. D. Garijo, S. Kinnings, L. Xie, L. Xie, Y. Zhang, P. E. Bourne, and Y. Gil, “Quantifying reproducibility in computational biology: the case of the tuberculosis drugome,” PloS one, vol. 8, no. 11, p. e80278, 2013.

58. S. Narayanasamy, Y. Jarosz, E. E. Muller, A. Heintz-Buschart, M. Herold, A. Kaysen, C. C. Laczny, N. Pinel, P. May, and P. Wilmes, “IMP: a pipeline for reproducible reference-independent integrated metagenomic and metatranscriptomic analyses,” Genome biology, vol. 17, no. 1, p. 260, 2016.

59. B. Kim, T. Ali, C. Lijeron, E. Afgan, and K. Krampis, “Bio-Docklets: virtualization containers for single-step execution of NGS pipelines,” GigaScience, 2017.

60. P. Dixon, “VEGAN, a package of R functions for community ecology,” Journal of Vegetation Science, vol. 14, no. 6, pp. 927–930, 2003.

61. E. Paradis, J. Claude, and K. Strimmer, “APE: analyses of phylogenetics and evolution in R language,” Bioinformatics, vol. 20, no. 2, pp. 289–290, 2004.

62. R. Suzuki and H. Shimodaira, “Pvclust: an R package for assessing the uncertainty in hierarchical clustering,” Bioinformatics, vol. 22, no. 12, pp. 1540–1542, 2006.

